# Constructing smaller genome graphs via string compression

**DOI:** 10.1101/2021.02.08.430279

**Authors:** Yutong Qiu, Carl Kingsford

## Abstract

The size of a genome graph — the space required to store the nodes, their labels and edges — affects the efficiency of operations performed on it. For example, the time complexity to align a sequence to a graph without a graph index depends on the total number of characters in the node labels and the number of edges in the graph. The size of the graph also affects the size of the graph index that is used to speed up the alignment. This raises the need for approaches to construct space-efficient genome graphs.

We point out similarities in the string encoding approaches of genome graphs and the external pointer macro (EPM) compression model. Supported by these similarities, we present a pair of linear-time algorithms that transform between genome graphs and EPM-compressed forms. We show that the algorithms result in an upper bound on the size of the genome graph constructed based on an optimal EPM compression. In addition to the transformation, we show that equivalent choices made by EPM compression algorithms may result in different sizes of genome graphs. To further optimize the size of the genome graph, we purpose the source assignment problem that optimizes over the equivalent choices during compression and introduce an ILP formulation that solves that problem optimally. As a proof-of-concept, we introduce RLZ-Graph, a genome graph constructed based on the relative Lempel-Ziv EPM compression algorithm. We show that using RLZ-Graph, across all human chromosomes, we are able to reduce the disk space to store a genome graph on average by 40.7% compared to colored de Bruijn graphs constructed by Bifrost under the default settings.

The RLZ-Graph software is available at https://github.com/Kingsford-Group/rlzgraph

## 1 Introduction

The linear reference genome suffers from reference bias that results in discarding informative reads sequenced from non-reference alleles during alignment [Ballouz et al., 2019]. To reduce the reference bias, alternative read alignment approaches that use a set of genomes as the reference have been recently introduced [Chen et al., 2020, Novak et al., 2017]. Genome graphs, due to their compact structure to store the shared regions of highly similar strings, are widely used to represent and analyze a collection of reference genomes compactly [Paten et al., 2017, Computational Pan-Genomics Consortium, 2018, Sherman and Salzberg, 2020, Sherman et al., 2019].

A genome graph of a collection of sequences is a labeled directed graph such that each sequence is equal to the concatenation of node labels on a path. We call such path a reconstruction path. The size of a genome graph is the space to store the graph structure, which is the set of nodes, edges and the node labels.

The size of a genome graph is crucial to the efficiency of operations such as mapping sequencing reads. Shown in Jain et al. [2019], the time complexity of mapping a string to a genome graph is directly correlated with the total number of characters in node labels and the number of edges. The speed of sequence-to-graph mapping can be further improved by a graph index, the size of which is also dependent on the size of the genome graphs. [Paten et al., 2017, Sirén et al., 2014, 2020].

Most of the existing genome graph construction algorithms do not directly optimize the size of the genome graph. Some of these algorithms choose a type of graph in order to adapt to a specific type of input data, such as read alignment [Garrison et al., 2018, Li et al., 2020, Paten et al., 2011, Mäkinen et al., 2020], variant calls [Garrison et al., 2018, Rakocevic et al., 2019, Dilthey et al., 2015] or raw sequencing reads [Iqbal et al., 2012], and then optimize the chosen graph. The other only optimize the graph index that stores reconstruction paths based an assumed type of genome graphs, for example, the variation graphs [Sirén et al., 2020, Sirén, 2017] or the colored compacted de Bruijn graphs [Almodaresi et al., 2017, 2020, Holley and Melsted, 2020, Muggli et al., 2019, Minkin et al., 2017]. As a result, the graphs constructed can be large in terms of both the space taken by the graph structure or the lengths of the reconstruction paths.

While a small genome graph is desirable, the smallest genome graph may be useless if each edge is allowed to be traversed multiple times. The smallest genome graph is a complete graph with four nodes, or *K*_4_, whose labels are *A, T, C* and *G*, respectively, and contains the reconstruction path for any genomic sequence. The strings stored in a genome graph are parsed into a sequence of nodes on the reconstruction path. However, the parsing of strings would have lengths equal to the lengths of the strings in *K*_4_, which undermines the goal of a genome graph to compactly store similar strings.

In order to construct a small genome graph that balances the size of the graph and the lengths of the reconstruction paths, we introduce the definition of a restricted genome graph and formalize the restricted genome graph optimization problem, which seeks to build a smallest restricted genome graph given a collection of strings.

We present a genome graph construction algorithm that directly addresses the restricted genome graph size optimization problem. Optimizing the size of a restricted genome graph is similar to optimizing the space taken by a set of strings, which echoes the external pointer macro (EPM) scheme. We introduce a pair of algorithms that transform between the EPM-compressed form and the restricted genome graphs, and prove an upper bound on the size of the restricted genome graph constructed given an optimized EPM-compressed form from a set of input sequences. We further reduce the number of nodes and edges in the constructed restricted genome graph by introducing and solving the source assignment problem via integer linear programming (ILP).

As a proof-of-concept that compression-based genome graph construction algorithms produce small genome graphs efficiently, we build the RLZ-Graph, which is based on an EPM scheme compression heuristic known as the relative Lempel-Ziv (RLZ) algorithm. The EPM compression problem is NP-complete [Storer and Szymanski, 1982]. Among the approximation heuristics to solve the EPM compression problem, the relative Lempel-Ziv algorithm [Kuruppu et al., 2010] runs in linear time and achieves good compression ratios on human genomic sequences [Deorowicz et al., 2015, Deorowicz and Grabowski, 2011, Ferrada et al., 2014].

We evaluate the performance of RLZ-Graph by comparing to the colored compacted de Bruijn graphs (ccdBG) [Iqbal et al., 2012]. CcdBG construction methods, similar to the compression-based genome graph construction algorithms, process the input sequences directly without intermediate steps such as alignment or variant calling. In ccdBG, the input sequences are fragmented into preliminary nodes that represent unique strings of length *k*, or *k*-mers. Each edge represents the adjacency between two *k*-mers in the sequences stored. The preliminary nodes with in-and out-degrees equal to 1 on a path are further merged into a supernode. Still, the number of nodes and edges, as well as the number of characters in node labels, in a ccdBG can increase significantly as the number of sequences stored increases. The size of the graph also depends heavily on the parameter *k*. These factors may offset the effort to efficiently encode the reconstruction path information in the graph indices [Almodaresi et al., 2017, 2020, Holley and Melsted, 2020, Muggli et al., 2019, Minkin et al., 2017]. Despite the different approaches to build the ccdBG indices, ccdBG construction methods results in the same underlying de Bruijn graph structure. When we compare our algorithm with ccdBG construction algorithms, we only compare the graph structure, which includes nodes, edges and sequences stored in each node.

To examine the performance of RLZ-Graph, we compare sizes of the RLZ-Graph with the ccdBG constructed by Bifrost [Holley and Melsted, 2020] on all human chromosome sequences from 100 individuals from the 1000 Genome Project [1000 Genomes Project Consortium, 2015]. The number of nodes and edges produced by RLZ-Graph are reduced significantly compared with the ccdBG. Across all chromosomes, the disk space taken to store the graph representation of 100 sequences is on average reduced by 40.7% compared with the ccdBG built under the default settings.

Additionally, we evaluate the performance of the ILP solution to the source assignment problem on RLZ-Graphs constructed from *E. coli* genome sequences, for which many whole genome sequences are available. We show that the solutions to the source assignment problem reduces the number of nodes by around 8% on 300 *E. coli* genomes.

## 2 Definitions

### 2.1 Strings

Let *s* be a string. *s*[*b*: *e*] denotes a substring starting from position *b* (inclusively) of *s* up to position *ε* (inclusively). We assume 0-indexing throughout this article. The length of *s* is denoted by |*s*|. Concatenations of strings {*s*_1_,…, *s_n_*} are denoted by *s′* = *s*_1_ · *s*_2_ ·… · *s_n_*.

### 2.2 Genome Graphs

#### Definition 1 (Genome graph)

*A genome graph G* = (*V, E, ℓ*) *of a collection of strings* 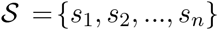 *is a directed graph with node set V, edge set E, and node labels £*(*u*) *for each node u. A genome graph of* 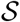 *contains a collection of paths* 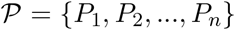, *where* 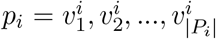, *such that* 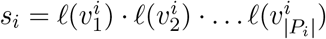 *for all* 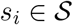. *Such paths are called reconstruction paths*.

The size of a genome graph *G* = (*V, E, ℓ*) is denoted by *size*(*G*), which is the space to store the set of nodes, edges and node labels (Section 3.1). The number of nodes in node set *V* and the number of edges in edge set *E* are denoted as |*V*| and |*E*|, respectively.

#### Definition 2 (Restricted genome graph)

*A restricted genome graph is a genome graph with a source and sink node and the restriction that each edge is allowed to be traversed at most once in all reconstruction paths*.

An example of a restricted genome graph is shown in Figure 1. Each edge is traversed only once in all reconstruction paths, and parallel edges are present. In a restricted genome graph, if we insert the source and sink nodes to the beginning and the end of each reconstruction path and add edges directing from sink to source, then the concatenation of reconstruction paths for all sequences forms an Eulerian tour. For a restricted genome graph *G* = (*V, E, ℓ*) and a collection of all reconstruction paths 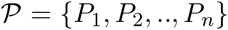, we have 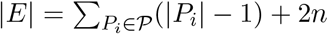, where 2*n* edges are the edges directing from source nodes and edges directing to sink nodes.

**Fig. 1.**
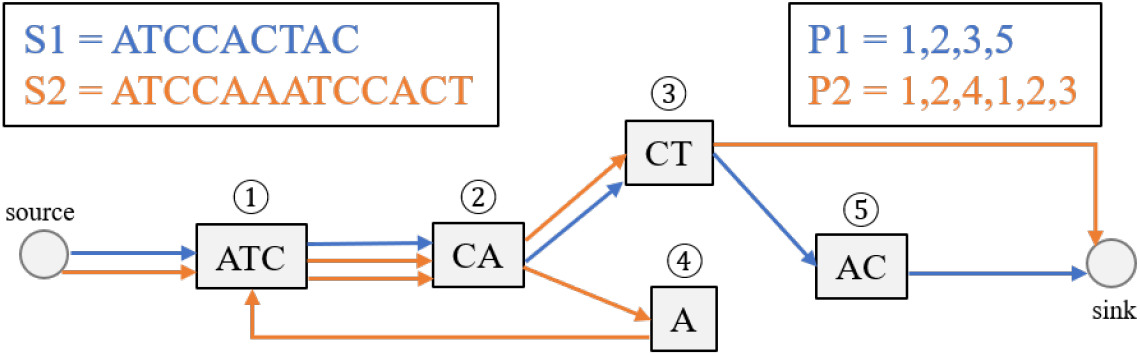
An example of a restricted genome graph. The graph stores two strings, S1 and S2. The color of the edges denotes the origin of node adjacencies.

### 2.3 External Pointer Macro (EPM) Compression Scheme

We review the definition of the external pointer macro (EPM) scheme for data compression [Storer and Szymanski, 1982].

#### Definition 3 (Pointers in EPM)

*Given a reference string R, a pointer p_i_* = (*pos_i_, len_i_*) *represents the substring R*[*pos_i_*,: *pos_i_, + len_i_* – 1].

We say that two pointers, *p_i_* = (*pos_i_, len_i_*) and *p_j_* = (*pos_j_, len_j_*) are equivalent to each other if *R*[*pos_i_*: *pos_i_*, + *len_i_* – 1] = *R*[*pos_j_*: *pos_j_* + *len_j_* – 1].

#### Definition 4 (External pointer macro (EPM) model)

*Given an alphabet Σ and a string T, a compressed form of string T adopts the EPM if the compressed data follows the form C* = *R*#*t*, where *R is a string over Σ, t* = *p*_1_*p*_2_,… *is a sequence of pointers that represent substrings in R*, # *is a separator symbol that is not in Σ, and T is equal to the string produced by substituting pointers in t by their corresponding substrings*.

The string *T* may represent a set of strings *S* = {*s*_1_, *s*_2_,…, *s_n_*} by concatenation, i.e. *T* = *s*_1_$*s*_2_…$*s_n_*, where $ ≠ # and $ ∈ *Σ*, where *Σ* is the alphabet for *S*. In this case, the compressed form *t* will also contain the character $ that separates sequences of pointers that represent different strings.

It is natural to consider optimizing the size of the compressed string *C*, *size*(*C*) (Section 3.2), which leads to:

#### Problem 1 (EPM decision problem [Storer and Szymanski, 1982])

Given a string *T* and an integer *m*, determine if there is a compressed form *C* of *T* that follows EPM such that *size*(*C*) ≤ *m*.

In Storer and Szymanski [1982], Problem 1 is shown to be NP-complete.

In EPM, the substring represented by a pointer may occur several times in the reference string.

We define such occurrences as sources of a pointer:

#### Definition 5 (Source)

*A source*, (*pos*_1_, *len*), *of a pointer p* = (*pos*_2_, *len*) *is an occurrence of R*[*pos*_2_: *pos*_2_ + *len* – 1] *in R*. *In other words, R*[*pos*_1_: *pos*_1_ + *len* – 1] = *R*[*pos*_2_: *pos*_2_ + *len* – 1]. *Each pointer p is associated with a source set* 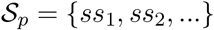, *where R*[*ss_i·pos_*: *ss_i·pos_* + *ss_i·len_* – 1] = *R*[*p·pos*: *p.pos* + *p.len* – 1] *for all* 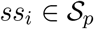.

Sources are used to refer to the occurrences of a substring on the reference string *R*, and pointers are used to refer to the pair of integers eventually stored in the compressed string *t*.

#### Definition 6 (Boundaries of sources and pointers)

*The boundaries of a source s* = (*pos, len*) *are defined as* (*b,ε*), *where b* = *pos and e* = *pos* + *len. b is the left boundary and e is the right boundary. The similar definition of boundaries applies to pointers*.

Two boundaries, (*b*_1_,*e*_1_) and (*b*_2_,*e*_2_), intersect if and only if *b*_1_ = *b*_2_ or *b*_1_ = *e*_2_ or *e*_1_ = *b*_2_ or *e*_1_ = *e*_2_.

### 2.4 Relative Lempel-Ziv Algorithm

#### Definition 7 (Right-maximal pointer)

*Given a reference string R and a string T, let pointer p* = (*pos, len*) *represent the substring R*[*pos*: *pos* + *len* – 1] *and T*[*pos*’: *pos*’ + *len* – 1]. *p is right-maximal if R*[*pos*: *pos* + *len*] ≠ *T*[*pos*’: *pos*’ + *len*].

#### Definition 8 (Phrase)

*Given a reference string R and a string T, a phrase, p* = (*pos, len*), *is a right-maximal pointer*.

The relative Lempel-Ziv (RLZ) algorithm, proposed by Kuruppu et al. [2010], runs in linear time and achieves good compression ratios with genomic sequences. RLZ takes a reference string *R* as input and parses the input string *T* greedily from left to right. At position *i* in *T*, RLZ substitutes the longest prefix of *T*[*i*: |*T*| – 1] that matches a substring in *R* with a phrase. Let the length of the phrase be *len*. After substitution, RLZ skips to position *i* + *len* in *T* and repeats the substitution process until *T* is exhausted. The process of phrase production is called RLZ factorization. In some analysis of RLZ, the reference string is generated from the set of input strings [Gagie et al., 2016]. Nevertheless, the RLZ factorization algorithm given a reference string remains the same.

The definitions introduced above are demonstrated in Figure 2, where *R* is the reference string and *T* is the input string to the RLZ algorithm. RLZ factors *T* into a sequence of three phrases, shown as *t*. The compressed form of the input string *T* is *C* = *R*#*t*. Each phrase is associated with some sources that are represented as line segments in the figure. For example, the last phrase, (7, 2), replaces the substring *T*[7: 8]. It also corresponds to two sources in R: (3, 2) and (7, 2), which are represented by the green line segments in R. The left and right boundaries of phrase (7,2) are (b = 7, e = 8) in *T*. Source (3,2) intersects with sources (1, 4) and (3,3). However, sources (1, 4) and (3, 3) do not intersect with each other.

**Fig. 2.**
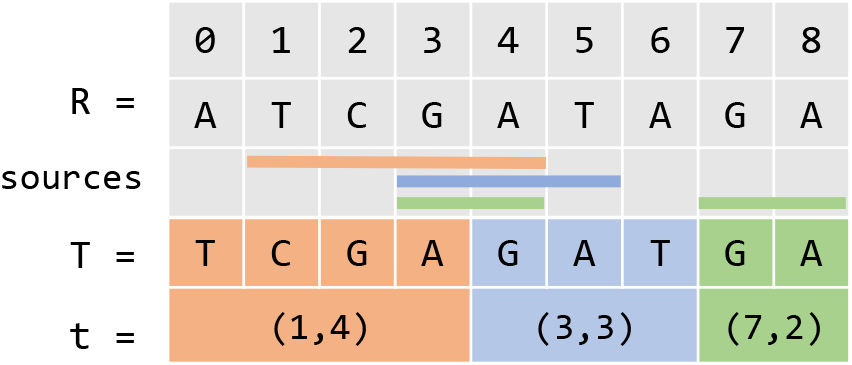
An example of RLZ factorization. The top row is the indices of characters in the strings. *R* is the reference string, *T* is the input string and *t* is a sequence of phrases resulted from RLZ factorization. Colored line segments on the third row represent the sources associated with phrases with the same color.

## 3 Size formulation of restricted genome graphs and EPM-compressed forms

### 3.1 Size of a genome graph

We adopt a natural formulation of the size of a labeled graph, which describes the space to store nodes, edges and the node labels. Given a restricted genome graph *G* = (*V, E,ℓ*) over alphabet *Σ*, let *L* be a string that contains every node label as a substring and *Σ* be the alphabet. Each node can be represented as a pointer to *L*, i.e. *v* = (*pos, len*), such that *ℓ*(*v*) = *L*[*pos*: *pos* + *len* – 1]. Each node takes 2 log |*L*| bits to store. The graph structure is stored as pairs of adjacent nodes. Each edge takes space 2 log |*V*| bits. Therefore, the total space taken by a restricted genome graph, denoted by *size*(*G*), under this model is:

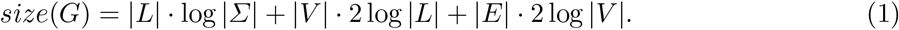

We introduce the restricted genome graph optimization problem:

#### Problem 2 (Restricted genome graph optimization problem)

Given a set of sequences, build a restricted genome graph *G* such that *size*(*G*) is minimized.

In the above formulation, note that |*E*| refers to the number of edges including the parallel edges. Solutions to Problem 2 avoid a trivial genome graph solution, that is a complete graph with four nodes, or *K*_4_, where each node has label *A, T, C*, and *G*, respectively. From *K*_4_, any sequence over the alphabet *Σ* = {*A, T, C, G*} can be reconstructed under the definition of a genome graph (Definition 1). However, the length of the reconstruction path would be equal to the length of the sequence, which undermines the purpose of the genome graph, part of which is to produce a short parsing of the input sequence.

In a restricted genome graph, the number of edges grows as the lengths of reconstruction paths increase. Therefore, minimizing the size of the restricted genome graph achieves a combined objective of a small genome graph and short parsing of the input sequences.

### 3.2 Size of an EPM-compressed form

Next, we consider the space taken by an EPM-compressed form *C* = *R*#*t*. The space taken by *C*, *size*(*C*), is the space to store the total number of unique pointers in *t*, the sequence *t* and the reference string *R*. We first encode each unique pointer with a pair of integers, (*pos, len*), which takes space 2 log |*R*| bits. If there are *n* unique pointers, *t* can be stored as a sequence of identifiers of the unique pointers using |*t*| log *n* bits. Therefore, the total space taken by an EPM-compressed form is

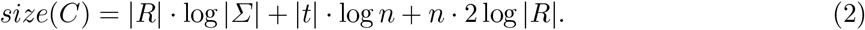

From equations (1) and (2), both the restricted genome graph and the EPM-compressed form have a size formulation that has three terms, which are the space taken by a reference string, the space taken by the unique pointers and the space to store the adjacencies between pointers.

In order to reduce the size of the restricted genome graphs (Definition 2), it is natural to borrow ideas from the field of string compression. We introduce two algorithms that transform between genome graphs and compressed strings produced by EPM compression scheme [Storer and Szymanski, 1982].

## 4 Transformation between EPM-compressed forms and genome graphs

### 4.1 EPM-compressed string to genome graph

Given an EPM-compressed form *C* = *R#t* of the original string *T*, and an alphabet *Σ*, the genome graph construction algorithm produces a restricted genome graph that stores both *R* and *T*.

A naïve algorithm to construct a genome graph is to create a node for each unique pointer in *t* and add an edge between nodes that represent each pair of pointers t[*i*] and t[*i* + 1]. However, in repetitive sequences such as the human genome, a substring may occur in several pointers and thus may be stored several times redundantly. In the example shown in Figure 3, the substring `AAA` would be stored three times according to the naäve algorithm, which results in excess space spent on storing repetitive content.

**Fig. 3.**
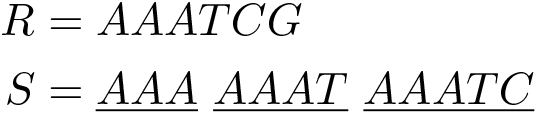
String *S* is factored into three pointers given the reference string *R*. Each underlined substring is represented by a different pointer. According to the naïve algorithm to construct the genome graph, three nodes are created from three pointers.

Our construction algorithm, introduced below as `two-pass CtoG`, merges the repetitive substrings shared by multiple pointers by grouping pointers by their positions on the reference. “Two-pass CtoG” creates nodes and edges of the genome graph in two passes through *t*. In the first pass, the algorithm creates nodes by cutting the reference string according to the boundaries of each pointer. In the second pass, the algorithm connects the nodes according to the adjacencies between pointers in the compressed string *t*.

#### First pass

Create a bit vector, 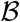. A bit set at 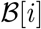 indicates that a pointer boundary falls at position *i* on *R*. Process *t* from left to right. For each pointer *p* = (*pos, len*), mark its boundaries in 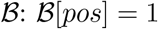 and 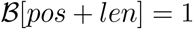. After *t* is exhausted, transform 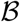 into an RRR data structure that supports rank operations in constant time [Raman et al., 2002], where 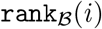 returns the number of set bits at or before position *i* in 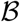. We then cut reference string at positions where a bit is set in 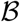. If 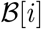 and 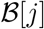 are the only set bits in the interval [*i*: *j*], we create a node *v* = (*pos, len*) = (*i,j* – *i*) with *ℓ*(*v*) = *R*[*i*: *j* – 1]. Each node can be treated as a pointer whose left and right boundaries are *i* and *j*, respectively. Each node is identified using its left boundary, i.e. 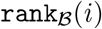.

We define the ordering of nodes. *V_i_* = (*pos_i_,len_i_*) ≺ *V_j_* = (*pos_j_,len_j_) iff *pos_i_*, < *pos_j_*, where *i* and *j* are the identifiers of *V_i_* and *V_j_*, and *i* < *j*. Add an edge between each *V_i_* and *v*_*i*+1_ for all *i* < |*V*| – 1. The path *v*_1_,*v*_2_,…, v*_|*V*|_ represents the reference string *R*.

#### Second pass

We process *t* from left to right again in the second pass. For each pair of pointers *t*[*i*] and *t*[*i* + 1], we need to connect the nodes that mark the right and left boundaries of *t*[*i*] and *t*[*j*], respectively. Let *t*[*i*] = (*pos_i_*, *len_i_*) and *t*[*i* + 1] = (*pos*_*i*+1_,*len*_*i*+1_). We need to find two nodes, *v_m_* = (*pos*_*m*_, *len*_*m*_) and *v_n_* = (*pos_n_*, *len_n_*), such that *pos_m_* + *len_m_* = *pos_i_*, + *len_i_* and *pos_n_* = *pos_i+1_*. Since each node is identified by their left boundary, two nodes can be identified by 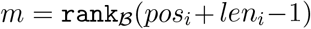 and 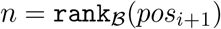. Edge (*v_m_, v_n_*) then represents the adjacency between *t*[*i*] and *t*[*i*+1] in *t*. Repeat the process until *t* is exhausted. Create a source and a sink node. Create an edge that connects the source to the first node of each compressed string and the last node of each compressed string to the sink. An example output of the algorithm is shown in Figure 4, where the compressed string is produced by the RLZ algorithm [Kuruppu et al., 2010].

**Fig. 4.**
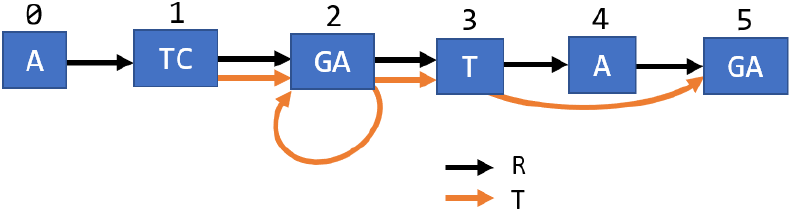
The RLZ-Graph of reference *R* = *ATCGATAGA* and input string *T* = *TCGAGATGA* (Figure 2). The black path 0,1,2,3,4, 5 encodes *R*, the orange path 1, 2,2,3, 5 encodes *T*. The parallel edges are shown for the purpose of illustration and are merged in the final graph.

The running time of the construction algorithm is *O*(|*t*| + |*R*|), where |*t*| is the total number of pointers. In both passes, the algorithm iterates through all pointers in *t* exactly once. Since the nodes are created by splitting the reference, there are at most |*R*| nodes and adding an edge between each pair of (*v_i_, v*_*i*+1_) takes *O*(|*R*|) time.

The constructed restricted genome graph stores the set of nodes, edges and the node labels. While storing the reconstruction paths is also important, it is a separate challenge from optimizing the graph structure. There has been a line of work that constructs small graph indices to store the reconstruction paths efficiently given any graph structure [Sirén et al., 2014, 2020, Sirén, 2017]. These indices can also be applied to our genome graph.

There are three types of edges in the produced restricted genome graph: the backward edges, the forward edges and the reference edges. We define the backward edges as edges that direct from *v_j_* to *v_i_*, where *j* ≥ *i*, which include self-loops. We define the forward edges as edges that direct from *v_i_* to *v_j_*, where *i* < *j* – 1. We define the reference edges as the edges that direct from *v_i_* to *v*_*i*+1_. In other words, reference edges (*v_i_* = (*pos_i_, len_i_*), *v_j_* = (*pos_j_, len_j_*)) connect nodes where the first node’s right boundary intersects with the second node’s left boundary, i.e. *pos_i_* + *len_i_* = *pos_j_*.

We show that the constructed graph is a restricted genome graph that contains reconstruction paths for *R* and *T* as in Theorem 1.

##### Theorem 1.

*Given an EPM-compressed form of string T, C* = *R#t, the algorithm described above creates a genome graph G* = (*V, E, ℓ*) *that contains reconstruction paths for R and T*.

*Proof*. In the second pass of the algorithm, edges are added between the nodes that are the suffix and the prefix of adjacent pointers. Therefore, all pointer adjacencies are represented as edges in the genome graph.

All substrings *R*[*i*: *j*] can be reconstructed from *G*. If *R*[*i*: *j*] is a substring of a node label, it can be reconstructed from *G*. If *R*[*i*: *j*] spans two nodes, it spans two nodes connected by a reference edge.

Any substring *T*[*i*: *j*] can be reconstructed from *G*. Suppose position *i* lands in the middle of a pointer *p_k_* = (*pos_k_, len_k_*), which means that *k* ≤ *i* ≤ *k* + *len_k_* – 1.

1. If *j* ≤ *k* + *len_k_* – 1, which means that *T*[*i*: *j*] is a substring of the string represented by a pointer. Since all pointers in *t* point to substrings in *R* and *R* can be reconstructed from *G*, a substring of a pointer can be reconstructed.
2. If *j* > *k* + *len_k_* – 1, which means that *T*[*i*: *j*] spans at least two pointers. From the previous case, we have that *T*[*i*: *k* + *len_k_* – 1] can be reconstructed using nodes and edges in *G*. Since all adjacencies between two pointers are represented in *G*, we can apply the analysis to the rest of *T*[*i*: *j*]. Therefore *T*[*i*: *j*] can be reconstructed if it spans more than one pointer.

### 4.2 Genome graph to EPM-compressed form

Given a restricted genome graph *G* = (*V, E, ℓ*) and a set of reconstruction paths 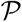 that represent strings in 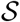, we present an algorithm, `GtoC`, that produces an EPM-compressed form *C* = *R#t* whose decompression equals string *T*, which is a concatenation of strings in 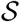.

Produce the reference string *R* by concatenating the node labels in an arbitrary order 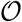. Each node can then be represented as a pointer to *R* and be denoted as *v_i_* = (*pos_i_, len_i_*), where *ℓ*(*v_i_*) = *R*[*pos_i_*: *pos_i_* + *len_i_* – 1]. Assign an identifier to each node such that for *v_i_* and *v_j_, i* < *j* if *pos_i_* < *pos_j_*.

Process all 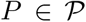 by substituting nodes with their pointer representations. If two adjacent nodes *v_i_* and *v_j_* in *P* are connected by a reference edge, merge the two nodes into one pointer *p* = (*pos_i_, len_i_*, + *len_j_*). Concatenate all processed *P*, which results in *t*. The converted sequence of pointers *t* is then *p*_1_, *p*_2_,…, *p*_|*t*|_, where 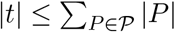.

The converted *C* satisfies the EPM definition where *R* is a string over *Σ* and *t* is a sequence of pointers to substrings in *R*. Since the concatenation of paths in 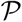 spells out *T* by concatenating all the labels of nodes on the path, substituting the pointers in *t* with corresponding substrings reconstructs *T*.

The running time of the construction algorithm is 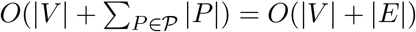.

The size of the produced EPM-compressed form can be further reduced if the reference string *R* is equal to the shortest superstring that contains all the node labels. However, finding the shortest superstring is a NP-hard problem when the number of nodes is greater than 2 and would be impractical in dealing with large genomes.

## 5 Upper-bound on the size of the restricted genome graph and the EPM-compressed form

We show that the size of a restricted genome graph *G* produced using the two-pass CtoG algorithm is bounded by the terms of the input EPM-compressed form *C* (Lemma 1).

### Lemma 1.

*Given an optimally compressed EPM form C* = *R#t, the size of the transformed restricted genome graph G* = (*V, E, ℓ*), *size*(*G*), *according `to two-pass CtoG` in Section 4.1 has an upper bound:*

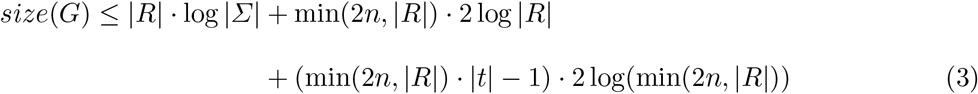

*where n is the number of unique pointers in t*.

*Proof*. The algorithm introduced in Section 4.1 creates nodes by cutting the reference string *R* according to boundaries of pointers. Each node is stored as a pointer (*pos, len*) to *R*, which takes 2 log | *R*| bits.

The total number of nodes produced by cutting the reference is ≤ min(2*n*, |*R*|). The number of cuts introduced by each unique pointer is ≤ 2. The maximum number of nodes given a reference string *R* is |*R*|. Therefore, the space to store all the nodes is ≤ min(2*n*, |*R*|) · 2 log |*R*|.

The total number of edges, including reference and non-reference edges, in a restricted genome graph is ≤ min(2*n*, |*R*|) · |*t*| – 1. After the first pass of `two-pass CtoG`, the interval corresponding to each pointer may be cut into several nodes. Let the average number of nodes contained in each pointer’s interval be *a* ≤ |*V*| ≤ min(2*n*, |*R*|). The average number of reference edges within each pointer is then *a* – 1. The total number of edges between pointers is |*t*| – 1, and the total number of edges within pointers is (*a* – 1) · |*t*|. Together, the number of edges in the reconstruction path is *a* · |*t*| – 1 ≤ min(2*n*, |*R*|) · |*t*| – 1.

The size of the genome graph is then:

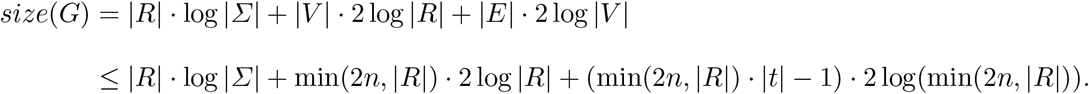

In practice, the graphs are stored such that the parallel edges are merged. We show that the size of the genome graph G’ produced by merging the parallel edges in *G* can also be bounded by the terms of the EPM-compressed form *C* (Lemma 2).

### Lemma 2.

*Given a restricted genome graph, G* = (*V,E,I*), *constructed from an optimally compressed EPM form C* = *R#t, the size of the genome graph, G′* = (*V, E′, ℓ*), *produced by merging parallel edges in G has an upper bound:*

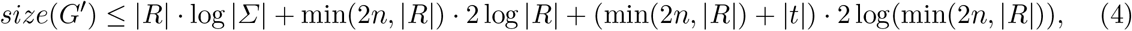

*where n is the number of unique pointers in t*.

*Proof*. Merging parallel edges does not change the number of nodes and the concatenation of node labels.

The number of reference edges in *G′* is equal to |*V*| – 1, as the nodes are produced by cutting the reference string.

The number of forward and backward edges in *G* is equal to |*t*| – 1, and the number of forward and backward edges in *G′* is ≤ |*t*| – 1 due to parallel edge merging. According to `two-pass CtoG`, since *C* is optimal, only a forward or a backward edge can be added for each pair of adjacent pointers in *t* during the second pass. Suppose two adjacent pointers, *p*_1_ = (*pos*_1_, *len*_1_) and *p*_2_ = (*pos*_2_, *len*_2_), result in a reference edge, which means that *pos*_2_ = *pos*_1_ + *len*_1_, the two pointers can be merged into *p*_3_ = (*pos*_1_, *len*_1_ + *len*_2_). Merging two pointers reduces the size of *C*, which contradicts the assumption that the size of *C* is optimal.

Together, the space to store all the edges in *G′* is ≤ |*V*| + |*t*| ≤ (min(2*n*, |*R*|)+|*t*|)·2 log min(2*n*, |*R*|).

Therefore, the size of the genome graph *G′* after merging the parallel edges in *G* is:

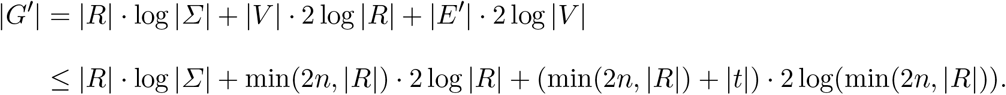

We show in Lemma 3 that the size of the EPM-compressed form produced by `GtoC` algorithm (Section 4.2) is upper-bounded by the terms of the size of a restricted genome graph.

### Lemma 3.

*Given a restricted genome graph G* = (*V, E, ℓ*) *of a collection of strings* 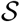, *the size of the transformed EPM-compressed form of the concatenated strings in* 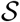, *C* = *R#t according to `GtoC` described in Section 4.2 has an upper bound:*

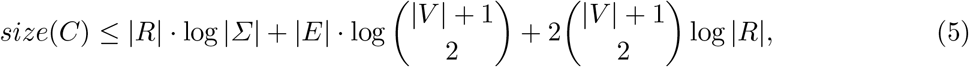

*where R is a string formed by concatenating all node labels*.

*Proof*. Let the set of paths corresponding to the set of strings 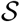 be 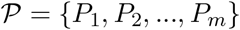, where *m* is the number of strings in 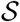. Since *G* has an optimal size, the number of edges in *G* is exactly 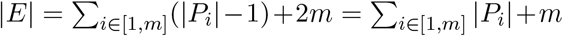, where *E* includes reference, forward and backward edges. Note that if an edge does not belong to any path in 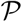, it can be eliminated in the graph, which results in a smaller restricted genome graph.

According to `GtoC`, the pointers are created by either directly converting a node or merging two nodes connected by a reference edge in a path 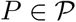. Let the number of reference edges be *r*. The number of pointers in *t*, or |*t*|, is equal to 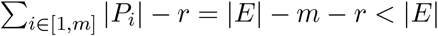.

Given |*V*| nodes, the reference constructed by concatenating all node labels contains |*V*| + 1 cut positions including the positions before *R*[0] and after *R*[|*R*| – 1]. From these cut positions, we can produce at most 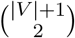 pointers by selecting two positions as boundaries of a pointer. Let the total number of unique pointers be *n*. Then 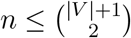.

Together, the size of the EPM-compressed form is:

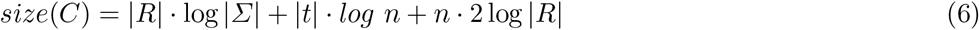

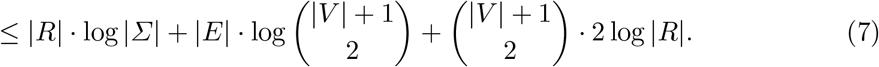

The pair of algorithms do not produce an optimal genome graph or optimal EPM-compressed form. Still, given an optimal input, the pair of algorithms achieve results that are bounded by the original terms in the input. We further improve the transformation from EPM-compressed form to genome graph by addressing the source assignment problem introduced below.

## 6 Source assignment problem

In an EPM-compressed form *C* = *R#t*, each pointer may be associated with a substring that occurs several times in *R*. We name such occurrences as sources. A source (*pos_i_, len_i_*) is assigned to a pointer *p* if *p* = (*pos_i_, len_i_*).

In the EPM formulation, assigning different sources to a pointer does not change the size of the compressed string. However, the assignment of sources may affect the number of nodes significantly. According to the `two-pass CtoG` algorithm, the number of cuts made in the reference is equal to the number of distinct pointer boundaries. Therefore, the choice of sources is directly related to the number of nodes in the graph. An example is illustrated in Figures 2 and 4. The last phrase, (7,2), is associated with two sources, (3,2) and (7,2). If we assign (3,2) to the phrase, which is different from the case in Figure 2, the number of nodes created will be 5. Otherwise, 6 nodes will be created as in Figure 4.

Given an EPM-compressed form and the set of sources corresponding to each pointer, if we can assign sources such that the total number of unique pointer boundaries is minimized, we can reduce the size of the created graph. We formulate the source assignment problem and present an integer linear programming (ILP) solution for the optimal source assignment in genome graph construction.

### Problem 3 (Source assignment problem)

Given a collection of sources sets 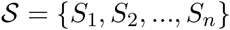, where *S_i_* denotes the set of sources for a unique pointer i, find a set of sources *S′* such that for all *S_i_*, 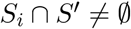 and 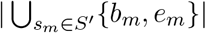 is minimized, where *b_m_, e_m_* are boundaries of source *m*.

In this problem, we choose one source for each pointer such that the union of boundaries {*b_m_, e_m_*} of each chosen source *s_m_* = (*pos_m_, len_m_*) is minimized. As a reminder, *b_m_* = *pos_m_* and *e_m_* = *pos_m_* + *len_m_*.

For convenience, we denote the union of boundaries in a source set *S* by 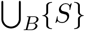, which is equivalent to 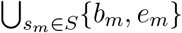.

The formulation of the source assignment problem is similar to the hitting set problem in that it chooses the minimum number of positions to hit every pointer. However, the objective is indirectly related to the number of the chosen sources, and the sources and pointers are defined in a string context. The hardness of the source assignment problem is open due to these differences from the setting of the hitting set problem. Still, the similarities to the hitting set problem lead to the formulation of an integer linear programming solution.

### 6.1 Integer linear programming formulation

The objective of the ILP is to minimize the number of cuts made in the reference, where each cut is made at the boundaries of chosen sources. For each chosen source *s* = (*pos_i_, len_i_*), a cut is placed at positions *pos_i_*, and *pos_i_*, + *len*, which are left and right boundaries of *s*.

We first construct a set of integers *I* that is the union of all source boundaries. Create a binary variable *x_p_* for each *p* ∈ *I. x_p_* is set to one if a cut is made at position *p*.

We create a binary variable *y_si_* for each source *s_i_* = (*pos_i_, len_i_*) that indicates whether the source is chosen. We create a constraint (Inequality 9) that at least one source is chosen from each set. We create another set of constraints (Inequalities 10,11) that ensures that if a source is chosen, two cuts are made at its left (*posf_i_* and right (*pos_i_*, + *len_i_*) boundaries. This leads to the ILP:

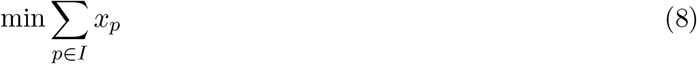

subjects to

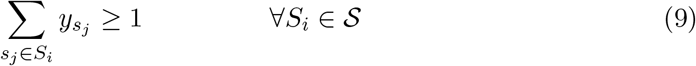

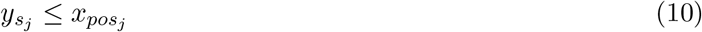

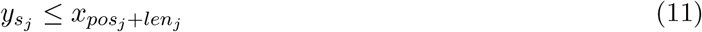

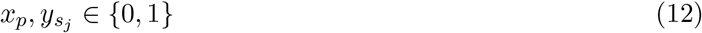

### 6.2 Pruning to reduce the number of sources

In practice, a pointer with a short length may correspond to a large number of sources. For example, a pointer with length one may correspond to |*R*|/4 sources, where *R* is the reference string when the alphabet size is 4. This could result in a huge number of variables in the ILP formulation and would hinder its practicality significantly.

To address this, we preprocess the sources as follows. If a source does not intersect with any other sources of different pointers, we eliminate the source from the source set unless it is the only source of a pointer. We name the eliminated sources isolated sources. Removing such sources does not affect the optimality of the solution.

#### Lemma 4.

*If a set of sources, S, that satisfies the constraints of the source assignment problem, includes an isolated source s, it is possible to find a set of sources S’ with equal or lower objective value that does not include s*.

*Proof*. Let the pointer for the isolated source be *p* and the source set of *p* be *S_p_*. Since *s* is an isolated source, there must be at least another source *s′* in *S_p_*. If *s′* also does not intersect with any other sources in *S*, 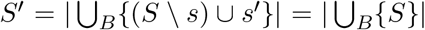. Otherwise, if *s′* intersects with some sources in *S*, this means that the union of source boundaries is reduced by at least 1 if we replace *s* with *s′*, i.e. 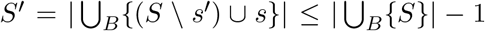. Therefore, excluding all isolating sources during preprocessing does not affect the optimality of the solution.

## 7 Relative Lempel-Ziv Graph

As a proof-of-concept that constructing a genome graph using a compression scheme results in small graphs, we implement the graph construction algorithm based on an EPM compression scheme algorithm, relative Lempel-Ziv. Given a reference string *R*, the relative Lempel-Ziv (RLZ) algorithm [Kuruppu et al., 2010], introduced in Section 2.4, seeks to greedily produce a compressed form of *R* where all pointers in *R* are right-maximal. We name the right-maximal pointers phrases. RLZ factorization in this manuscript is done on the compressed suffix array in the SDSL C++ library [Gog et al., 2014].

We apply the `two-pass CtoG` algorithm described in Section 4.1 to construct a RLZ-Graph. We merge the parallel edges in the implementation as it is the common practice in genome graph storage.

An example of RLZ-Graph is shown in Figure 4. The RLZ-Graph is constructed based on the RLZ factorization in Figure 2, where the reference string is *R* = ATCGATAGA, the input string is *T* = *TCGAGATGA* and the factored phrase sequence is *t* = (1,4), (3,3), (7,2). The nodes are produced by segmenting *R* according to the boundaries of sources assigned to phrases in *t*.

In the implementation of RLZ-Graph, we build a bi-directed graph where each node can be traversed in forward and reverse directions, which is a commonly applied technique in other genome graph construction algorithms. For each node *v* = (*pos, len*), *pos* is referred to as the head of the node and *pos* + *len* is referred to as the tail. If a node is traversed in reverse direction, its label is denoted as 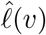, which is equal to the reverse complement of *ℓ*(*v*). This technique is useful in genomic sequences that underwent structural variations such as inversions, where the entire genomic segment is replaced by its reverse complement due to a double-strand break. During the construction of the RLZ-Graph, we use a modified reference sequence *R* by concatenating the reference genome of the organism of interest with its reverse complement. Before the source assignment step, we mark each source as reversed if it is located on the reverse complement half of *R* and translate its boundary positions to the forward half. After the source assignment step, we mark a pointer as reversed if it is assigned a reversed source. When we add edges, if we encounter a reverse pointer *p* = (*pos, len*), we add an edge directing to the tail of the node *v_i_* = (*pos_i_, len_i_*) and an edge directing from the head of the node *v_j_* = (*pos_j_, len_j_*), where *pos_i_* = *pos* and *pos_j_* + *len_j_* = *pos* + *len*.

We ran all our experiments on a server with 24 cores (48 threads) of two Intel Xeon E5 2690 v3 @ 2.60GHz and 377 GB of memory. The system was running Ubuntu 18.04 with Linux kernel 4.15.0.

### 7.1 Performance of RLZ-Graph compared to the colored compacted de Bruijn graphs

We compare the size of the colored compacted de Bruijn graphs [Iqbal et al., 2012] with that of RLZ-Graphs on human genomic sequences. While there have been many graph construction algorithms for building colored de Bruijn graphs, the graph structure of ccdBG remains the same in these algorithms despite the different approaches to store the reconstruction paths as identifiers in each node. The comparisons made in this section only concern the graph structure, which includes the nodes, edges and the node labels.

## 8 Experimental results

We use Bifrost [Holley and Melsted, 2020] to construct the ccdBG. The genome graphs constructed include nodes, labels of nodes and edges, are stored in graphical fragment assembly (GFA) format [Li, 2016]. In GFA file, the nodes of a graph are stored as a list of pairs of node identifiers and labels, and edges are stored as a list of pairs of node identifiers. Same as the RLZ-Graph, the graph constructed by Bifrost is bi-directed and does not contain parallel edges. The RLZ-Graph produced in this section does not use the ILP solution to assign sources due to the time and memory concern. Instead, we adopt the leftmost heuristic, where the leftmost source is assigned to each pointer. A source *s_i_* = (*pos_i_*, *len_i_*) is to the left of source *s_j_* = (*pos_j_*, *len_j_*) if *pos_i_* < *pos_j_*.

We build the graphs on all human chromosomes and show the results on chromosome 1 here (see Supplementary Figure S1–S3 for the rest of the chromosomes). The genomes we use are from the 1000 Genome Project phase 3 [1000 Genomes Project Consortium, 2015]. For each chromosome, we randomly choose 5, 25, 50, 75 and 100 samples and generate their genomic sequences using the consensus command from bcftools [Li, 2011]. We construct both graphs using the sample sequences and the reference hg37. Hg37 is also used as the reference string in RLZ factorization. We vary the *k*-mer sizes used for Bifrost and report the sizes of graphs with *k* = 31, 63 and 127. The default choice of *k* of Bifrost is 31. We repeat each experiment 5 times.

Shown in Figure 5, we compare the graph size in different aspects. From 5 sequences up to 100 sequences, the graph produced by RLZ-Graph is smaller than the graph produced by Bifrost with different choices of *k* under all measures in the figure. At 100 sequences, the GFA file that stores the RLZ-Graph is 37% smaller than the GFA file storing the colored de Bruijn graph produced by Bifrost with *k* = 63 and is 42.2% smaller when *k* = 31 (Figure 5(d)). The number of total characters in the concatenated node labels are constant in the RLZ-Graph regardless of the increase of the number of sequences because nodes are produced by cutting a reference string.

**Fig. 5.**
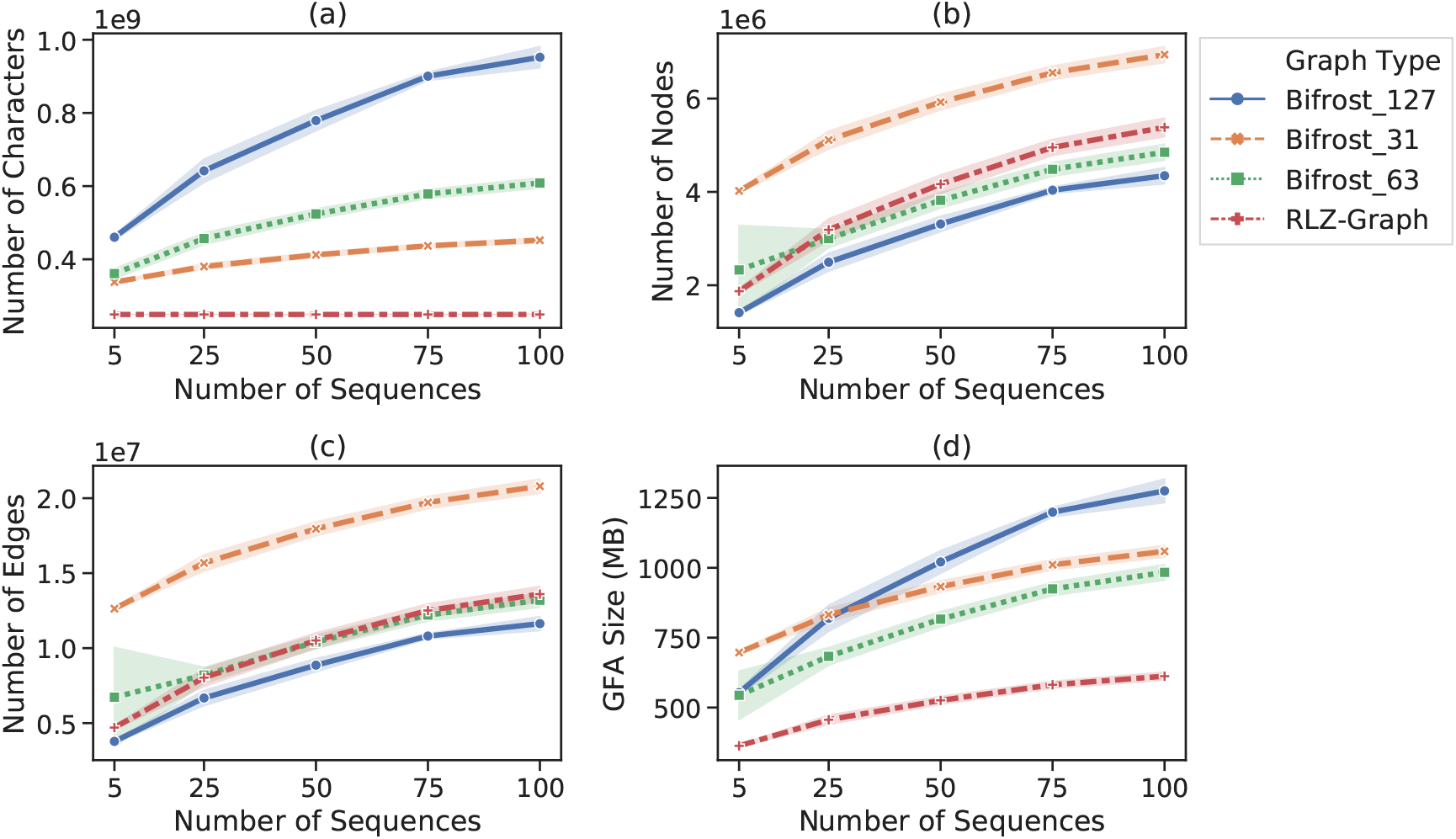
Comparison between RLZ-Graph and ccDBG constructed by Bifrost with *k* = 31, 63 and 127 on human chromosome 1 sequences. (a) Total number of characters in the node labels. (b) Number of nodes. (c) Number of edges. (d) Size of the GFA file that stores the graph structure and node labels.

The average running time of RLZ-Graph and Bifrost with *k* = 31, 63 and 127 on chromosome 1 is reported in Table 1. It takes RLZ-Graph around 2.5 hours to build a graph with 100 chromosome 1 sequences. The running time includes the time to do RLZ factorization. In all experiments, Bifrost is run in parallel in 20 threads while RLZ-Graph is run in a single thread. The RLZ-Graph implementation is not optimized and not parallelized compared to implementation of Bifrost. Still, the running time is on the similar scale compared to Bifrost.

**Table 1.** Average wall-clock running time of RLZ-Graph and Bifrost with different *k* values on chromosome 1 sequences.

| Number of sequences | 5 | 25 | 50 | 75 | 100 |
| --- | --- | --- | --- | --- | --- |
| **RLZ-Graph time (s)** | 1031 | 2707 | 4758 | 6872 | 9136 |
| **Bifrost k=31 time (s)** | 396 | 656 | 986 | 1542 | 1852 |
| **Bifrost k=63 time (s)** | 280 | 510 | 793 | 1126 | 1332 |
| **Bifrost k=127 time (s)** | 412 | 733 | 1098 | 1443 | 1744 |

When *k* = 15, the size of the GFA file that stores the ccdBG is 15 gigabytes for 5 chromosome 1 sequences and the running time is around 8 hours. Both the size of the graph and the running time is impractical compared to other *k* values. When *k* = 3, the size of the GFA file is 4.2 kilobytes for 5 chromosome 1 sequences with 32 nodes and 127 edges and the running time is around 2.5 hours. Although the graph is small, it is similar to the *K_4_* solution to the genome graph size optimization problem, where the length of the reconstruction path is approximately the same as the original string. With small *k*-mers, the ccdBG becomes impractical for human chromosomes. Therefore, we do not include the results of graphs constructed with smaller *k* values in Figure 5.

### 8.1 Performance of various source assignment heuristics

Aside from the ILP solution to the source assignment problem, sources are chosen by other heuristics in literature regarding RLZ factorization [Kuruppu et al., 2011]. Specifically, the leftmost source on the reference string is chosen (Left), or the lexicographically smallest source is chosen (Lex). The lexicographical order of sources of the same source set is defined such that *s_i_* = (*pos_i_*, *len_i_*) < *s_j_* = (*pos_j_*, *len_j_*) if *R*[*pos_i_*: |*R*| – 1] < *R*[*pos_j_*: |*R*| – 1] given a reference string *R*. In our implementation, a phrase is assigned to its lexicographically smallest source by default. In this section, we compare the performance of different source assignment heuristics in terms of the number of RLZ-Graph nodes in Figure 6(c).

**Fig. 6.**
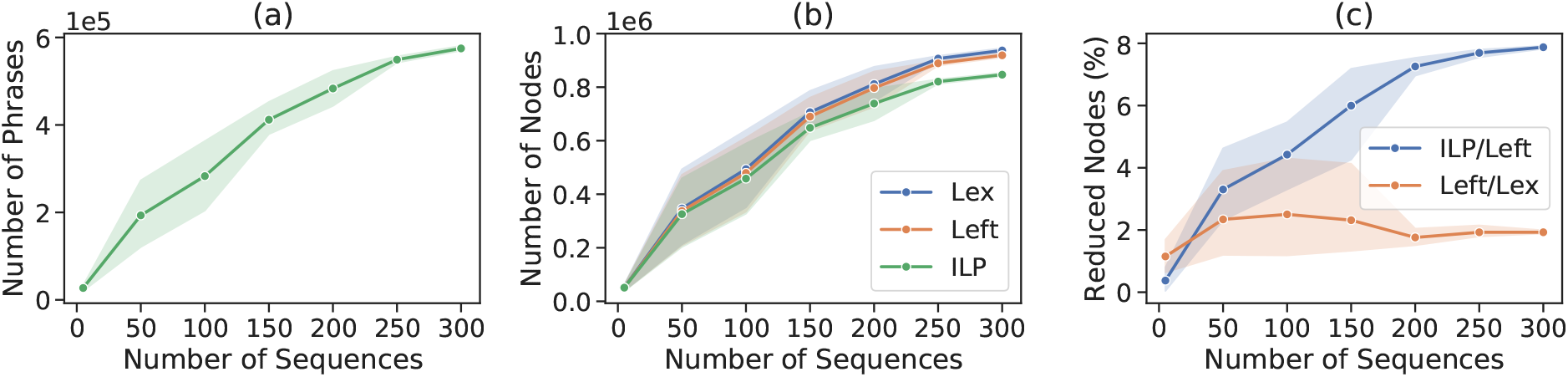
Performance of heuristics solving the source assignment problem. (a) The number of phrases. (b) The number of nodes. (c) Percentage of nodes reduced using the leftmost heuristic and the ILP solution during the source assignment step. The shaded area in the plots represents the standard deviation across 5 experiments and each data point in the figure represents the mean across 5 experiments. Lex: lexicographical heuristic. Left: leftmost heuristic. ILP: ILP solution.

We obtain 300 genomic sequences of *E. coli* O157 strain from Genbank [Clark et al., 2016], from which we randomly permute the genomic sequences and construct the RLZ-Graph on varying number of sequences. The first sequence in the randomly permuted 300 sequences is used as the reference string. We repeat each experiment 5 times.

In Figure 6(a), we show the rate at which the number of phrases produced by the RLZ factorization increases as the number of sequences increases. In Figure 6(b), we show the number of nodes produced due to different source assignment strategies. The ILP solution has the best performance and results in the fewest nodes. The percentage of reduced nodes is around 8% for 300 *E. coli* sequences. As the number of sequences increases, the ILP solution is able to eliminate more nodes compared to the heuristic that always chooses the leftmost source. The percentage of reduced nodes is calculated as 1 – (|*V*|_*ILP*_/|*V*|_*Left*_) and 1 – (|*V*|_*Left*_/|*V*|_*Lex*_), respectively.

## 9 Discussion

We define the restricted genome graph and formalize the restricted genome graph size optimization problem. The optimization problem balances both the size of the graph structure and the length of the reconstruction paths of sequences stored in the graph, which is similar to the string compression problem. Inspired by the similarity, we present a pair of algorithms that bridge genome graph construction and the external pointer macro model. We prove an upper bound on the size of the genome graph that is constructed based on an optimal compressed string from the EPM model. One key advantage of our graph construction algorithm is that the total number of characters stored in the graph remains the constant regardless of the number of sequences stored in the graph, which helps reduce the space taken by a genome graph. Further, since the number of nodes and edges are derived from an already compressed representation of strings, the number of nodes and the number of edges remain small.

We show that equivalent choices made by data compression algorithms may affect the size of the genome graph differently. To address this challenge, we introduce the source assignment problem, and present an ILP solution to it. We show that solving the source assignment problem prior to graph construction reduces the number of nodes by around 8%. Although it is a relatively small percentage, when dealing with very large genome graphs, it translates into substantial space saving. The application of solutions to the source assignment problem is not limited to the relative Lempel-Ziv algorithm, but can be applied to any EPM-compressed form to reduce the number of nodes and edges. The NP-completeness of the source assignment problem is still open.

As a proof-of-concept that compressed-based genome graph construction algorithms can produce small genome graphs, we implement RLZ-Graph based on the relative Lempel-Ziv algorithm [Kuruppu et al., 2010]. We show that using RLZ-Graph, we are able to reduce the size of the graph significantly on disk compared to the colored compacted de Bruijn graph. The choice of *k*-mer sizes is important in de Bruijn graph construction as it significantly affects the size of the graph. RLZ-Graph removes this dependence on the choice of *k* and produces practical graphs with a smaller size, which is scalable to the entire human genome. Although the implementation of RLZ-Graph is not optimized, its running time is comparable to the parallelized and optimized ccdBG construction method, Bifrost [Holley and Melsted, 2020]. A future direction would be to improve the implementation of RLZ-Graph by parallelizing the RLZ factorization step.

This work is an initial investigation into the connection between genome graph construction and string compression. We show that using compression algorithms, we can build small genome graphs efficiently, which opens up the possibilities in future research in adapting other data compression schemes to genome graph construction.

## Funding

This work has been supported in part by the Gordon and Betty Moore Foundation’s Data-Driven Discovery Initiative through Grant GBMF4554 to C.K., by the US National Institutes of Health (R01GM122935), and the US National Science Foundation (DBI-1937540). Y.Q. was supported by the Carnegie Mellon University School of Computer Science Cancer Research Fellowship for 2021.

## Financial Disclosure

C.K. is a co-founder of Ocean Genomics, Inc.

## Supplementary results

### 1 Comparison between ccdBGs and RLZ-Graphs on human chromosomes 2–22

We compare the sizes of genome graphs constructed on human chromosomes 2–22 by RLZ-Graph and Bifrost in Figures S1-S3. The experiment settings are the same as in Section 7.1 of the main text. Bifrost is run with *k* = 31, which is the default setting.

**Fig. S1.**
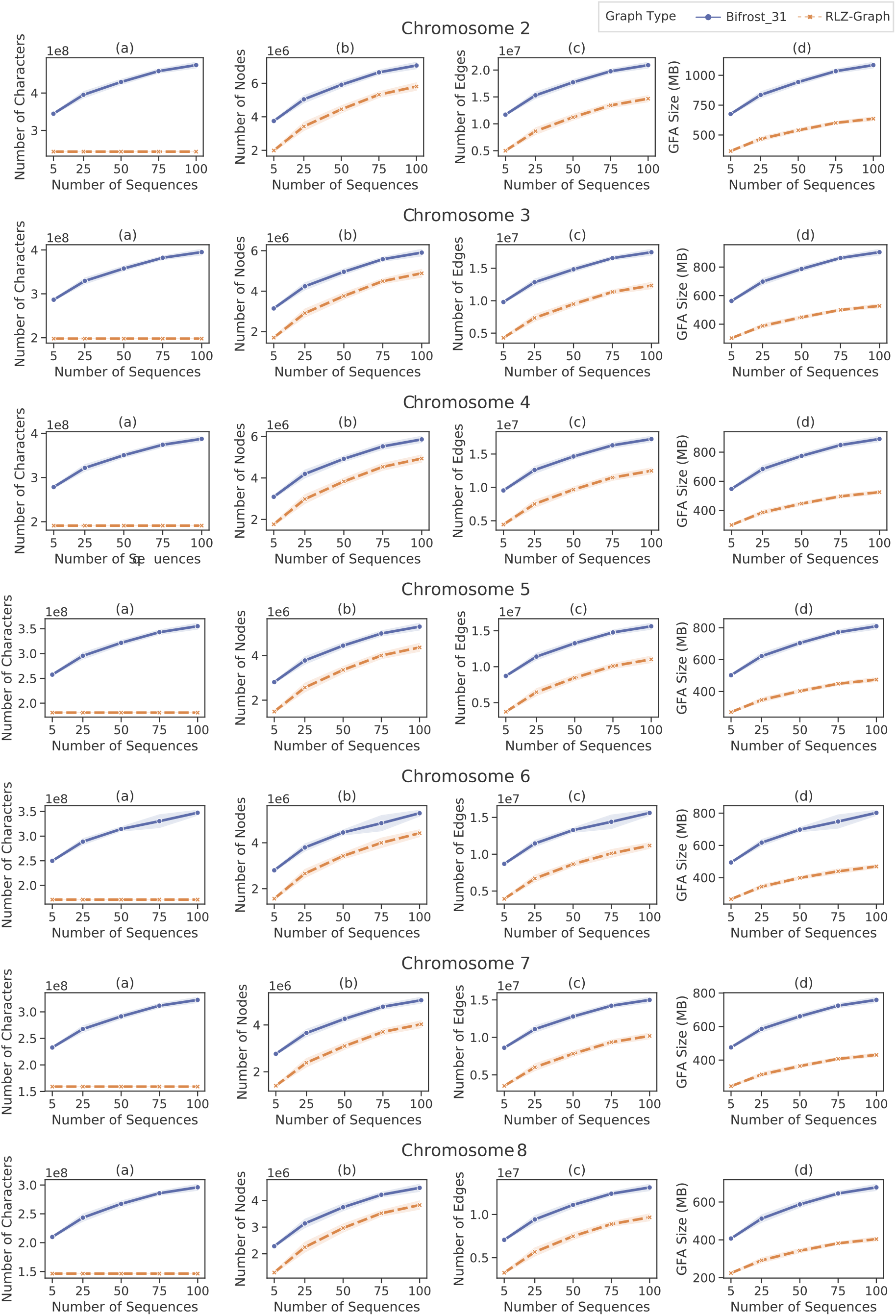
Comparison between RLZ-Graph and ccdBG constructed by Bifrost with *k* = 31 on human chromosomes 2–8. (a) Total number of characters in the node labels. (b) Number of nodes. (c) Number of edges. (d) Size of GFA file that stores the graph structure and node labels.

**Fig. S2.**
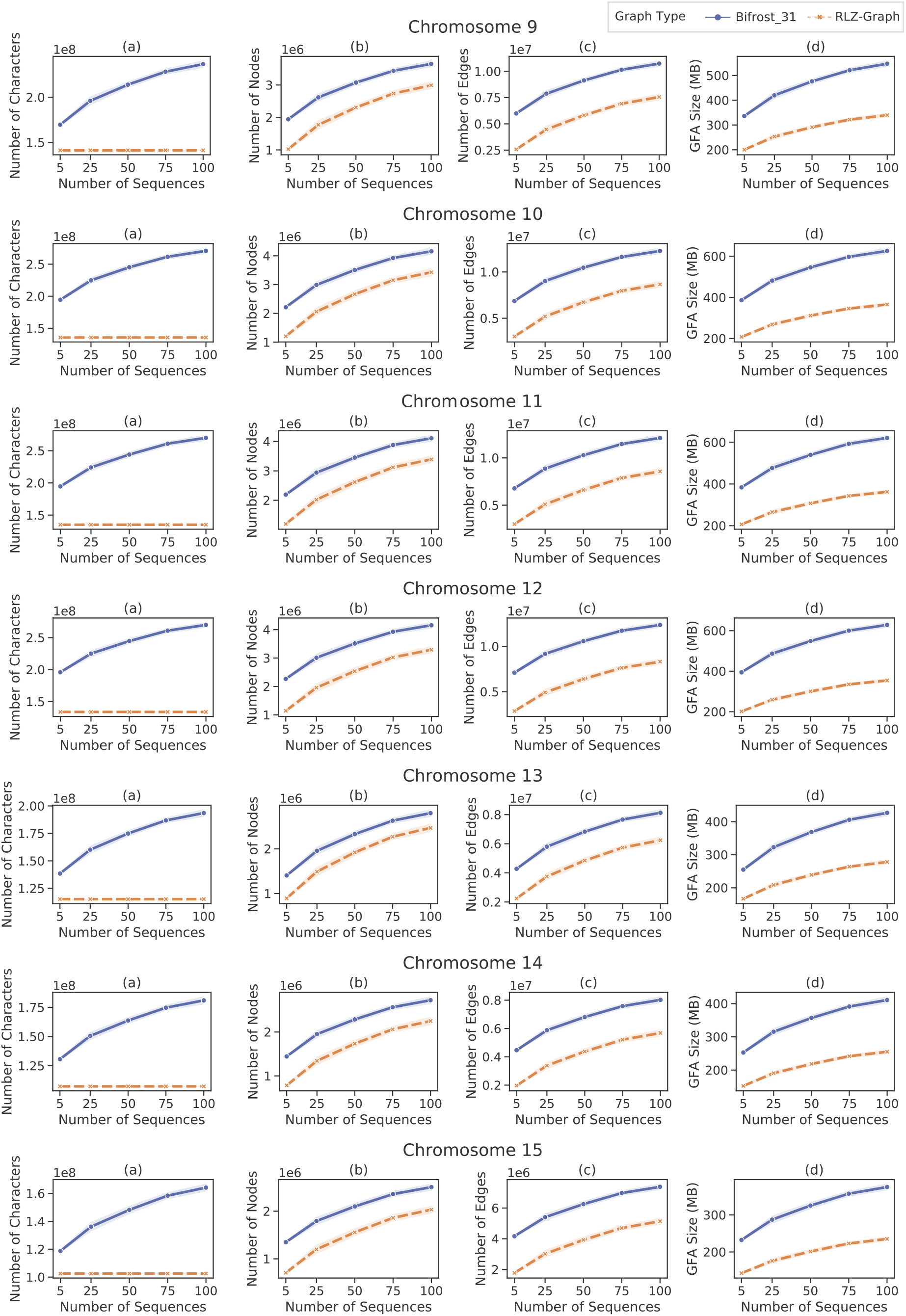
Comparison between RLZ-Graph and ccdBG constructed by Bifrost with *k* = 31 on human chromosomes 9–15. (a) Total number of characters in the node labels. (b) Number of nodes. (c) Number of edges. (d) Size of GFA file that stores the graph structure and node labels.

**Fig. S3.**
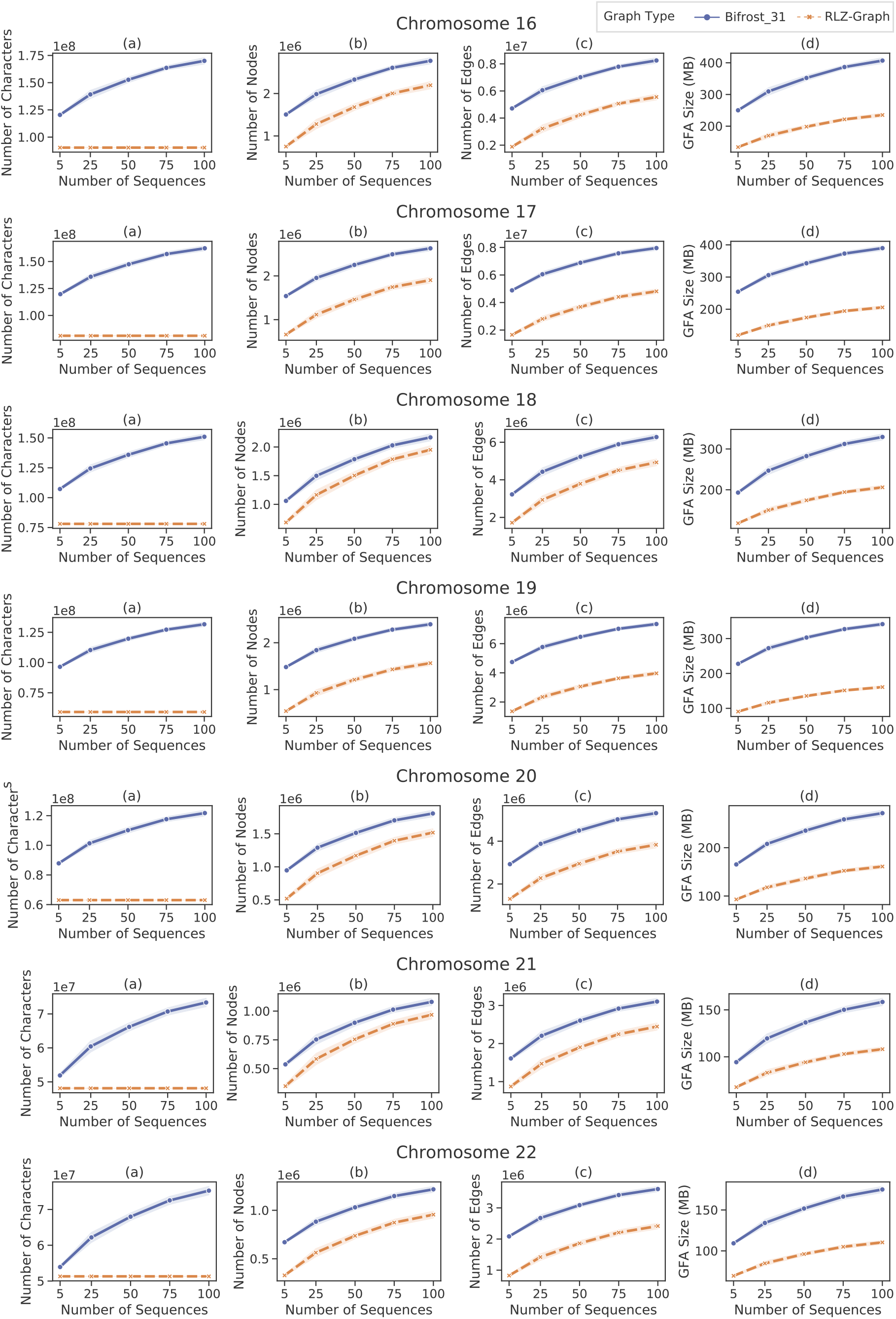
Comparison between RLZ-Graph and ccdBG constructed by Bifrost with *k* = 31 on human chromosomes 16–22. (a) Total number of characters in the node labels. (b) Number of nodes. (c) Number of edges. (d) Size of GFA file that stores the graph structure and node labels.

## Notes

### Competing Interest Statement

Carl Kingsford is a co-founder of Ocean Genomics, Inc.

